# Inhibition of Hsp70 suppresses neuronal hyperexcitability and attenuates seizures by enhancing A-type potassium currents

**DOI:** 10.1101/256602

**Authors:** Fang Hu, Jingheng Zhou, Yanxin Lu, Lizhao Guan, Ning-ning Wei, Zhuo Huang, Yi-Quan Tang, KeWei Wang

**Affiliations:** Qingdao University School of Phamacy, Qingdao 266021, China; Department of Molecular and Cellular Pharmacology, Peking University School of Pharmaceutical Sciences, Beijing 100191, China

**Keywords:** epilepsy, Hsp70, hyperexcitability, Kv4, KChIP4a

## Abstract

The heat shock protein 70 (Hsp70) is upregulated in response to stress and has been implicated as a stress marker in temporal lobe epilepsy (TLE). However, whether Hsp70 plays a pathologic or protective role in TLE remains unclear. Here we report that Hsp70 exerts an unexpected deleterious role in kainic acid (KA)-induced seizures, and inhibition of Hsp70 suppresses neuronal hyperexcitability and attenuates both acute and chronic seizures via enhancing A-type potassium currents primarily formed by Kv4 α-subunits and auxiliary KChIPs. Proteosomal degradation of Kv4-KChIP4a channel complexes is enhanced by Hsp70, which can be reversed by the Hsp70 inhibitors, 2-phenylethynesulfonamide (PES) and VER-155008 (VER). In cultured hippocampal neurons, either PES or VER can increase A-type Kv4 current to suppress neuronal hyperexcitability. Mechanistically, Hsp70-CHIP complexes directly bind to the N-terminus of auxiliary KChIP4a and target Kv4-KChIP4a complexes to the proteasome. Our findings reveal a previously unrecognized role of Hsp70 in mediating degradation of Kv4-KChIP4a complexes and regulating neuronal excitability, thus highlighting a therapeutic potential for hyperexcitability-related neurological disorders through Hsp70 inhibition.

## INTRODUCTION

Epilepsy is a common neurological disorder that afflicts over 65 million people worldwide and is characterized by spontaneous recurrent seizure activity (Chang & Lowenstein, 2003). Anti-epileptic drugs available over the past few decades are only effective in subsets of patients, and developing a better treatment and an eventual cure of epilepsy requires deeper understanding of pathogenic events and molecular changes that drive the aberrant neuronal firing (Loscher et al., 2013).

Heat shock proteins (HSP) are ubiquitous molecular chaperones that play important roles in a variety of biological processes ranging from cellular stress response to trafficking and quality control of membrane receptors (Hartl et al., 2011). In the mammalian brain, the predominant HSPs (Hsp70 and Hsp90) facilitate protein folding and guide proteins for ubiquitin-mediated degradation through interactions with co-chaperones such as CHIP and BAG-1 (Luders et al., 2000; Meacham et al., 2001). Chaperone-based therapies have been shown to be effective for nervous system diseases. Activation/overexpression of Hsp70 serves a protective role in various neurodegenerative diseases such as the polyQ diseases (Adachi et al., 2003; Kazemi-Esfarjani & Benzer, 2000; Wang et al., 2013), amyotrophic lateral sclerosis (ALS) (Bruening et al., 1999; Kieran et al., 2004) and Parkinson disease (PD) (Auluck et al., 2002; Klucken et al., 2004). Hsp70 upregulation, however, is not always beneficial. It is reported that inhibition of Hsp70 reduces tau levels in models of tauopathy and attenuates synaptic plasticity deficits, whereas activating Hsp70 stabilizes tau proteins both in cells and brain tissues (Abisambra et al., 2013; Jinwal et al., 2009). In various autoimmune diseases such as multiple sclerosis (MS) and experimental autoimmune encephalomyelitis (EAE), upregulation of Hsp70 can exacerbate the chronic inflammatory environment and promote myelin autoantigen recognition (Asea et al., 2000; Asea et al., 2002; Mycko et al., 2008), suggesting a complex role of Hsp70 in neurological diseases. Hsp70 has been suggested to be a stress marker and its expression is upregulated during status epilepticus (Yang et al., 2008). Although previous studies have shown that exogenous Hsp70 is able to attenuate seizure severity and improve neuronal survival (Ekimova et al., 2010; Yenari et al., 1998), the role of endogenous Hsp70 in the pathogenesis of epilepsy remains unclear.

Kv4 α-subunits co-assemble with the cytosolic Kv channel-interacting proteins (KChIPs) and the dipeptidyl peptidase-like proteins (DPLPs) to encode A-type potassium currents that play a critical role in regulating somatodendritic excitability (An et al., 2000; Jerng & Pfaffinger, 2014; Nadal et al., 2003; Soh & Goldstein, 2008; Zagha et al., 2005). A number of studies indicate that neuronal hyperexcitability in epilepsy is associated with changes in Kv4 channels (Bernard et al., 2004; Gross et al., 2016; Lugo et al., 2008; Monaghan et al., 2008; Singh et al., 2006; Tsaur et al., 1992). Auxiliary KChIP1-4 consist of a conserved C-terminal core domain with four EF-hand-like calcium binding motifs, and an alternatively spliced variable N-terminal domain that mediates diverse modulation on Kv4 function (Burgoyne, 2007). We recently reported that a specific splicing variant KChIP4a, distinct from other KChIPs, suppresses Kv4 surface expression via its unique N-terminal Kv4 channel inhibitory domain (KID) (Tang et al., 2013; Tang et al., 2014). Despite the unique modulation of Kv4 by KChIP4a, the pathophysiological significance of KChIP4a in central nervous system and neurological disorders is still unknown.

In the present study, we show an unexpected deleterious role of Hsp70 in kainic acid (KA) model of temporal lobe epilepsy (TLE) in rats. Pharmacological inhibition of Hsp70 reduces neuronal firing in primary cultured hippocampal neurons and suppresses KA induced acute and chronic seizures by enhancing A-type Kv4 currents. Furthermore, Hsp70 induced by epileptic stress reduces Kv4 expression through binding to the N-terminal KID of auxiliary KChIP4a subunits and facilitating proteasomal degradation of Kv4-KChIP4a complexes. Our findings highlight an important role of Hsp70 in regulating neuronal excitability, and inhibition of Hsp70 may lead to a therapeutic strategy for epilepsy or other hyperexcitability-related neurological disorders.

## RESULTS

### Hsp70 inhibition attenuates KA-induced acute seizures

To investigate the role of Hsp70 in epilepsy, we started examining the time course of Hsp70 expression in rat hippocampus of KA kindled epilepsy. Status epilepticus (SE) occurred several minutes after intracerebroventricular (ICV) injection of 1 μg KA, displaying electrographic seizures of high-frequency, high-voltage, rhythmic activity with clear onset and termination, and prominent voltage spikes preceding ictal events as seen in the hippocampal electroencephalogram (EEG) (Fig. 1A). SE lasted for at least 24-48 h after KA injection, and disappeared completely at 72 h (Fig. S1A). Western blot analysis showed that Hsp70 protein expression was significantly increased at 12 h, 24 h and 48 h after KA injection and returned to normal at 72 h (Fig. 1B), which was coincident with induction of SE in the rat KA model (Fig. S1A). In contrast, Hsp70 protein expression is not altered in the saline-injected control hippocampus (Fig. S1B). The Hsp70 mRNA was also dramatically increased in hippocampus 24 h after KA injection (Fig. S1C). These results indicate a long-lasting activation of Hsp70 during the development of epilepsy.

**Fig. 1.**
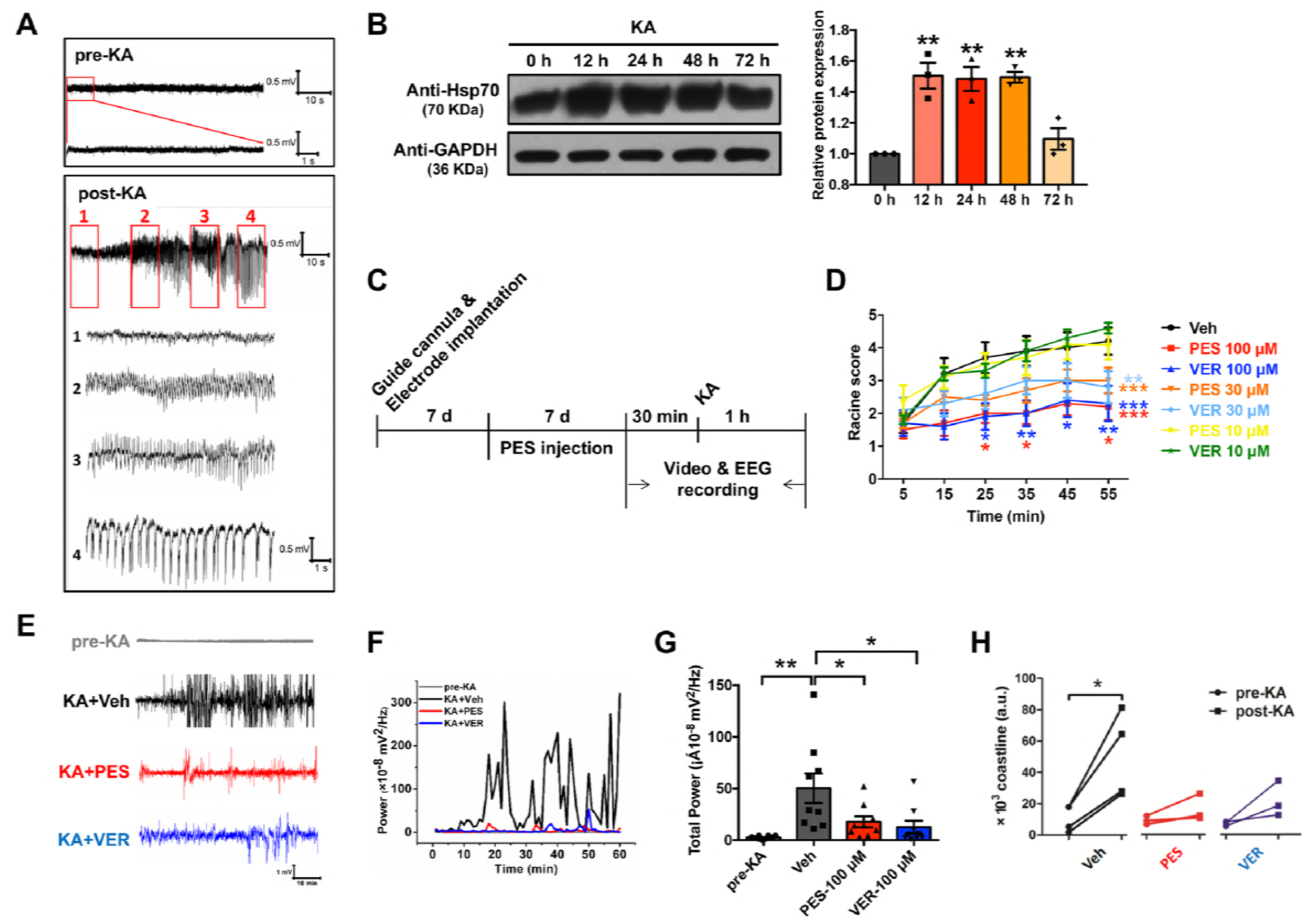
Concentration-dependent suppression of KA-induced acute seizures byHsp70 inhibitors in rats. **A** Representative EEG traces prior to (baseline) and after intracerebroventricular injection of 1 μg kainic acid (KA) in rat. Numbers and frames indicate regions of the seizures shown below at higher resolution. **B** Western blot analysis of Hsp70 from hippocampal lysates of KA-treated rats at indicated times. Data were normalized to 0 h; n = 3 independent experiments. **C** Experimental protocol for Hsp70 inhibitors test in acute seizure models. **D** Development of seizure behaviors after ICV administration of 1 μg KA in rats treated with vehicle, PES or VER at different concentrations (10, 30 or 100 μM, dissolved in 1 μl ACSF). Seizures were scored on the Racine Scale as described;n = 10. **E** Representative EEG traces of rats from pre-KA, KA + Veh (vehicle), KA + PES (100 μM), and KA + VER (100 μM) group. **F** The representative plots of power analyses of EEG using a similar timescale inpanel E. **G** EEG total power counts of rats from pre-KA, KA + Veh, KA + PES (100 μM),and KA + VER (100 μM) group. n = 9. Pre-KA in panel E, F and G represented before KA injection in KA + Veh group. **H** Coastline index counts of rats from Veh, PES (100 μM), and VER (100 μM)group; n = 3-4. All data are expressed as the means ± s.e.m.; comparisons were analyzed using one-way ANOVA followed by Dunnet’s tests (compared each group with 0 h or Veh) for B and G (excluding pre-KA group), unpaired Student’s *t* test for G (pre-KA and Veh group), two-way ANOVA (different time points) followed by Bonferroni’s multiple comparisons tests (compared each group with Veh) for D and paired Student’s *t* test in each group (compared post-KA with pre-KA) for H, *\*p* <0.05; *\*\*p*<0.01.

Overexpression of Hsp70 during the development of SE suggests its involvement in the pathogenesis of SE. To test this hypothesis, we assessed the effect of Hsp70 inhibitors on epileptic behaviors quantified in Racine’s scale (Racine, 1972), and seizure activities recorded by EEG in KA induced models of epilepsy. 2-phenylethynesulfonamide (PES) and VER-155008 (VER) are two recently identified small molecule inhibitors of Hsp70, targeting different sites in Hsp70 (Leu et al., 2009; Williamson et al., 2009). Rats were pre-treated with different doses of PES/VER (10 μM, 30 μM and 100 μM) once a day for 7 days before ICV administration of 1 μg KA (Fig. 1C). Both PES and VER showed a dose-dependent anticonvulsant effect (Fig. 1D). We also analyzed the total EEG power as a surrogate marker to assess the anti-seizure effect of Hsp70 inhibitors. There was no difference in EEG between vehicle control and Hsp70 inhibitor-treated groups before KA injection, indicating that repeated injections do not disrupt the circuits (Fig. S1D).

Quantitative analysis of the EEG power revealed that rats injected with KA displayed larger total power (50.3 ± 14.3 x 10^-8^ mV^2^/Hz) than before KA injection (2.5 ± 0.4 x 10^-8^ mV^2^/Hz) (Fig. 1E-G). In contrast, pre-administration of either PES (100 μM, 17.8 ± 5.3 x 10^-8^ mV^2^/Hz) or VER (100 μM, 12.5 ± 6.2 x 10^-8^ mV^2^/Hz) significantly reduced the total EEG power (Fig. 1E-G), indicating the anti-seizure effect of Hsp70 inhibition. Coastline parameter as an index for the cumulative difference in EEG (White et al., 2006) was significantly increased after KA administration in vehicle (Veh) group, but not in PES or VER treated group (Fig. 1H). As a control, treatment of animals with PES or VER produced no significant differences in spontaneous locomotor activity (Fig. S1E). These *in vivo* results demonstrate the anticonvulsant effects of Hsp70 inhibitors on KA-induced acute seizures in rats.

### Hsp70 inhibition attenuates KA-induced chronic seizures

To further investigate the anticonvulsant effects of Hsp70 on chronic seizures we generated a rat model of epilepsy that features spontaneous recurrent seizures (SRSs). 2-3 weeks after KA administration, SRSs developed and rats were injected with PES (100 μM) once a day for 7 days. In the vehicle control group rats received 1 μl ACSF (containing 2% DMSO) (Fig. 2A). We found that the administration of PES dramatically reduced the occurrence of epileptic seizures from a baseline of 7.4 ± 1.3 per day to 2.3 ± 0.62 per day, a mean decrease of 69% (Fig. 2B-C). In contrast, the seizure frequency in the vehicle group slightly increased after DMSO injection (Fig. 2B-C). We also examined electroencephalographic seizure activities before and after PES treatment. There were no obvious epileptiform discharges, such as spikes or sharp waves before the development of SRSs (Fig. 2D and S1A). After development of SRSs, the epileptiform discharges appeared almost every day (Fig. 2D). In contrast, PES treatment significantly decreased the total time of epileptiform discharges per day in each rat recorded (Fig. 2E), indicating that Hsp70 inhibitor exhibits an anticonvulsant effect on behavioral seizures and epileptiform activities in rat chronic spontaneous epilepsy. Collectively, our data strongly show that the stress protein Hsp70 is involved in epilepsy and inhibition of Hsp70 attenuates chronic seizures induced by KA in rats.

**Fig. 2.**
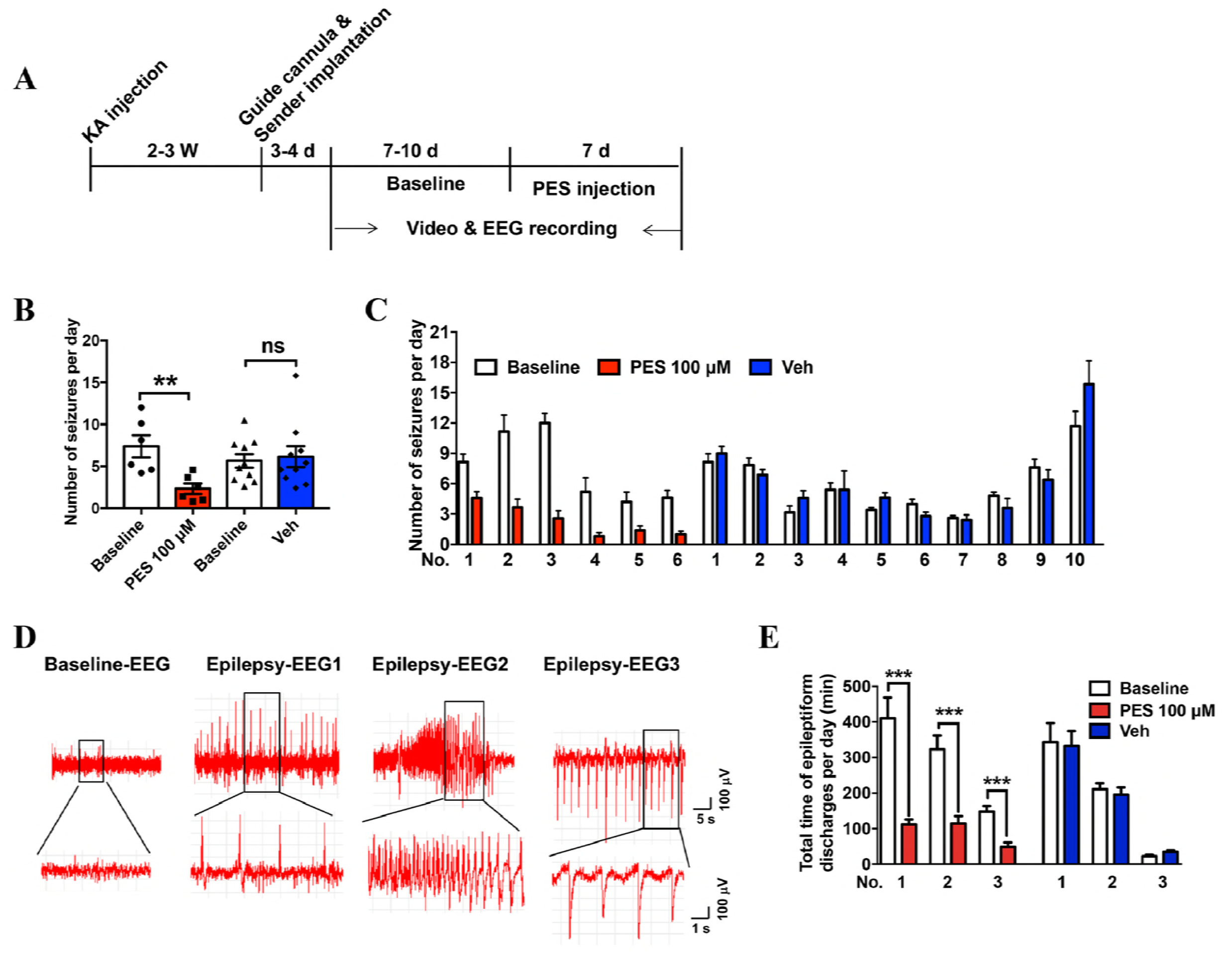
Inhibition of Hsp70 suppresses KA-induced chronic seizures in rats. **A** Illustration of the experimental procedures. **B** Statistical analysis for the number of seizures per day. Data were from three individual experiments with 10 rats in the Veh group and 6 rats in the PES group;comparisons were analyzed using paired Student’s *t* test in each group, \*\**p* <0.01. **C** Number of seizures per day of individual animals in Veh and PES group. **D** Representative EEG traces during normal station (baseline) and spontaneous seizures respectively. Frames indicate regions of the EEG traces shown below at higher resolution. **E** Statistical analysis of the total time of epileptiform discharges per day. Data were from three individual rats in the Veh group and PES group; comparisons were analyzed using paired Student’s *t* test in each group, \*\*\**p* <0.001.

### Suppression of neuronal excitability and enhancement of A-type current by inhibition of Hsp70

Since both Hsp70 inhibitors PES and VER suppressed KA-induced seizures, we tested whether inhibition of Hsp70 could regulate neuronal excitability. Neuronal firing was recorded by injecting 1000-ms depolarizing currents with different intensity into primary cultured hippocampal neurons pre-incubated with either PES (100 μM) or VER (100 μM) for 4 h. PES and VER-treated neurons produced significantly fewer action potentials, as compared with control neurons (Fig. 3A-B). The average time to first spike onset in response to 30 pA current injection was increased from 37.3 ± 3.5 ms in control neurons (n = 40) to 63.4 ± 11.8 ms (n = 19) in PES- or 65.3 ± 9.7 ms (n = 25) in VER-treated neurons (Fig. 3C). Action potential threshold and resting membrane potential were not affected by the treatment of PES or VER (Fig. 3D-E). We also recorded the neuronal firing of hippocampal neurons after incubation with KA. The results showed that pre-incubation of KA for 24 h or 0.5 h (data not shown) had no obvious effect in hippocampal neuronal excitability (Fig. S2A-B), likely due to the lack of enough synaptic connections and synchronized neuronal network activity in primary cultured hippocampal neurons.

**Fig. 3.**
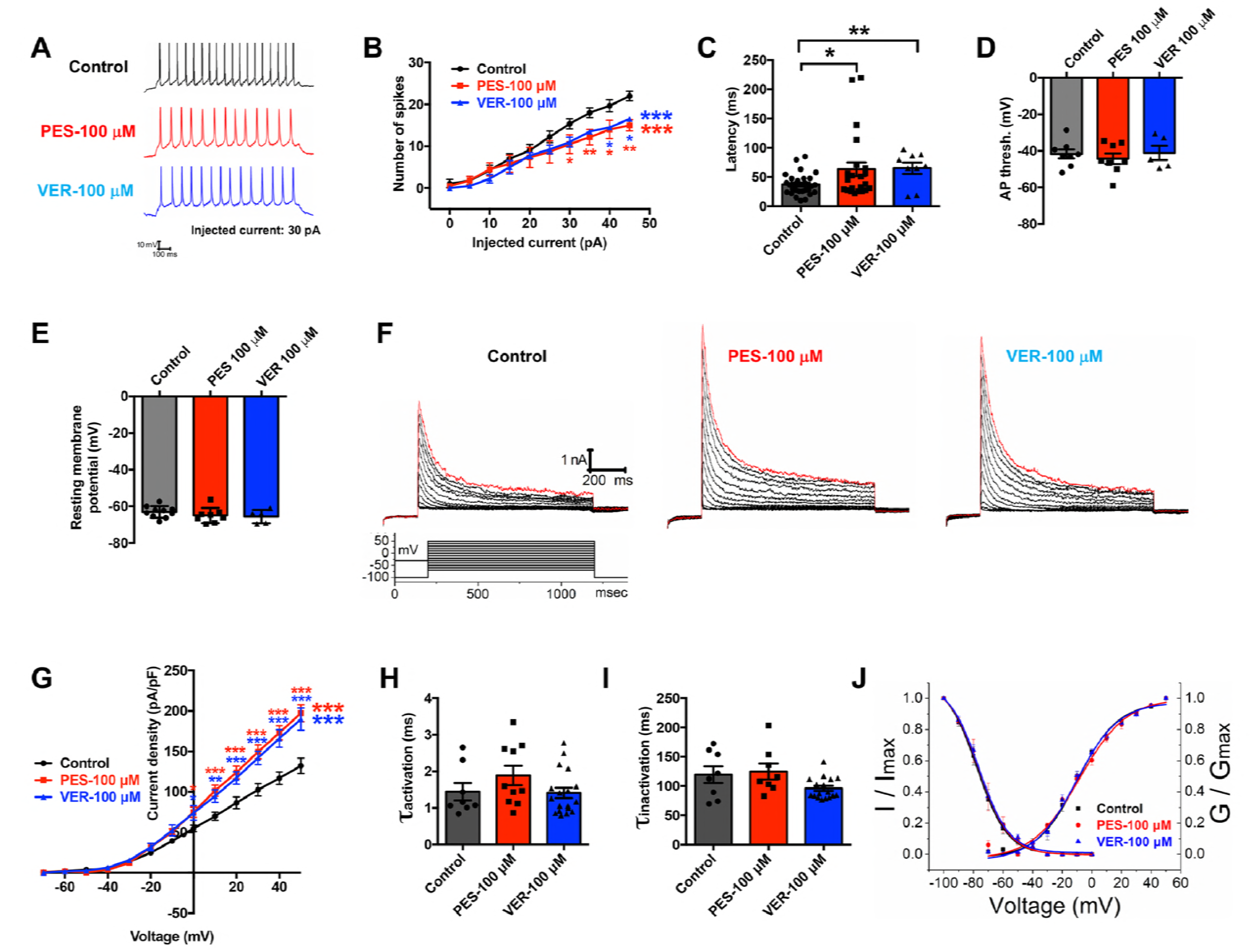
Reduction of neuronal excitability and enhancement of A-type current by Hsp70 inhibition. **A** Rat hippocampal neuron spiking activity was evoked by a squared depolarizing current pulse (30 pA for 1000 ms) in the absence or presence of PES and VER. **B** A summary of statistical analysis for firing frequency from panel A; n = 19-40 neurons. **C** A summary of statistical analysis for action potential latency in response to 30 pA current injection, n = 19-40 neurons. **D** A summary of statistical analysis for action potential threshold; n = 5-8 neurons. **E** A summary of statistical analysis for resting membrane potential; n = 5-10 neurons. **F***I_A_* recorded from primary cultured hippocampal neurons in the absence or presence of PES (100 μM) or VER (100 μM). **G**Current densities were plotted against membrane potentials; n = 10-11 neurons. **H**Quantitative analysis of activation time constants for *I_A_* in primary cultured hippocampal neurons in the presence or absence of PES or VER (100 μM). n =10-11 neurons. **I**Quantitative analysis of inactivation time constants for *I_SA_* in primary cultured hippocampal neurons in the presence or absence of PES or VER (100 μM). n = 10-11 neurons. **J** The voltage dependence of activation and inactivation of *I_A_* in primary cultured hippocampal neurons in the presence of PES or VER (100 μM). n = 9-11neurons. All values are expressed as the means ± s.e.m.; Comparisons for difference were analyzed using two-way ANOVA followed by Bonferroni’s multiple comparisons tests (compared each group with Control) for B and G, and one-way ANOVA followed by Dunnet’s tests (each group compared with Veh) for C-E, H and I, \**p* <0.05; \*\**p* <0.01; \*\*\**p* <0.001.

It was previously reported that increased excitability of CA1 pyramidal neuron dendrites in TLE is resulted from an activity-dependent reduction of A-type potassium current (*I*_*A*_) (Bernard et al., 2004) and the first spike latency is increased in hippocampal neurons when *I_A_* is increased (Kim et al., 2005). Thus, we hypothesized that Hsp70 inhibitors PES or VER might affect the *I_A_*. To test this hypothesis, we recorded *I_A_* using the prepulse protocol (Shibata et al., 2000) by subtracting the current evoked by test pulse after a 1000 ms voltage step at −30 mV (prepulse) at which *I_A_* is completely inactivated, from the current evoked from −100 mV (prepulse) (Fig. S2C). The subtracted current could be completely blocked by 4-aminopyridine (4-AP, 5 mM), indicating that the isolated current was *I_A_* (Fig. S2D). Incubation with PES (100 μM) or VER (100 μM) for 4 h dramatically increased the isolated *I_A_* currents (Fig. 3F-G), but had no significant effects on gating properties of *I_A_*. Neither activation and inactivation kinetics, nor the voltage dependence of activation and steady-state inactivation of *I_A_*, was altered by PES and VER (Fig. 3H-J). These data demonstrate that inhibition of Hsp70 enhances *I_A_* and reduces neuronal excitability.

### Hsp70 inhibition reduces Kv4 degradation caused by KA-induced seizures

*I_A_* in hippocampal and cortical neurons is encoded by Kv4.2 or Kv4.3 α-subunits (Norris & Nerbonne, 2010). To determine if KA-induced epilepsy or Hsp70 inhibition affected Kv4.2 or Kv4.3 currents, we measured the expression level of Kv4.2 and Kv4.3 α-subunits in rat hippocampus of KA-induced seizure models. The results showed that expression levels of both Kv4.2 and Kv4.3 α-subunits were significantly decreased in the hippocampi of KA-kindled rats and remained low up to 72 h after KA injection (Fig. 4A). Kv4.2 and Kv4.3 protein levels remained unchanged in the saline injected control group (Fig. S3). In contrast, inhibition of Hsp70 by PES (100 μM) or VER (100 μM) dramatically increased the expression of Kv4.2 and Kv4.3 α-subunits in rat hippocampi from both KA-induced acute and chronic seizure models (Fig. 4B-C). These results show that downregulation of Kv4.2 and Kv4.3 proteins during seizures is mediated by Hsp70 induction, which can be reversed by Hsp70 inhibition.

**Fig. 4.**
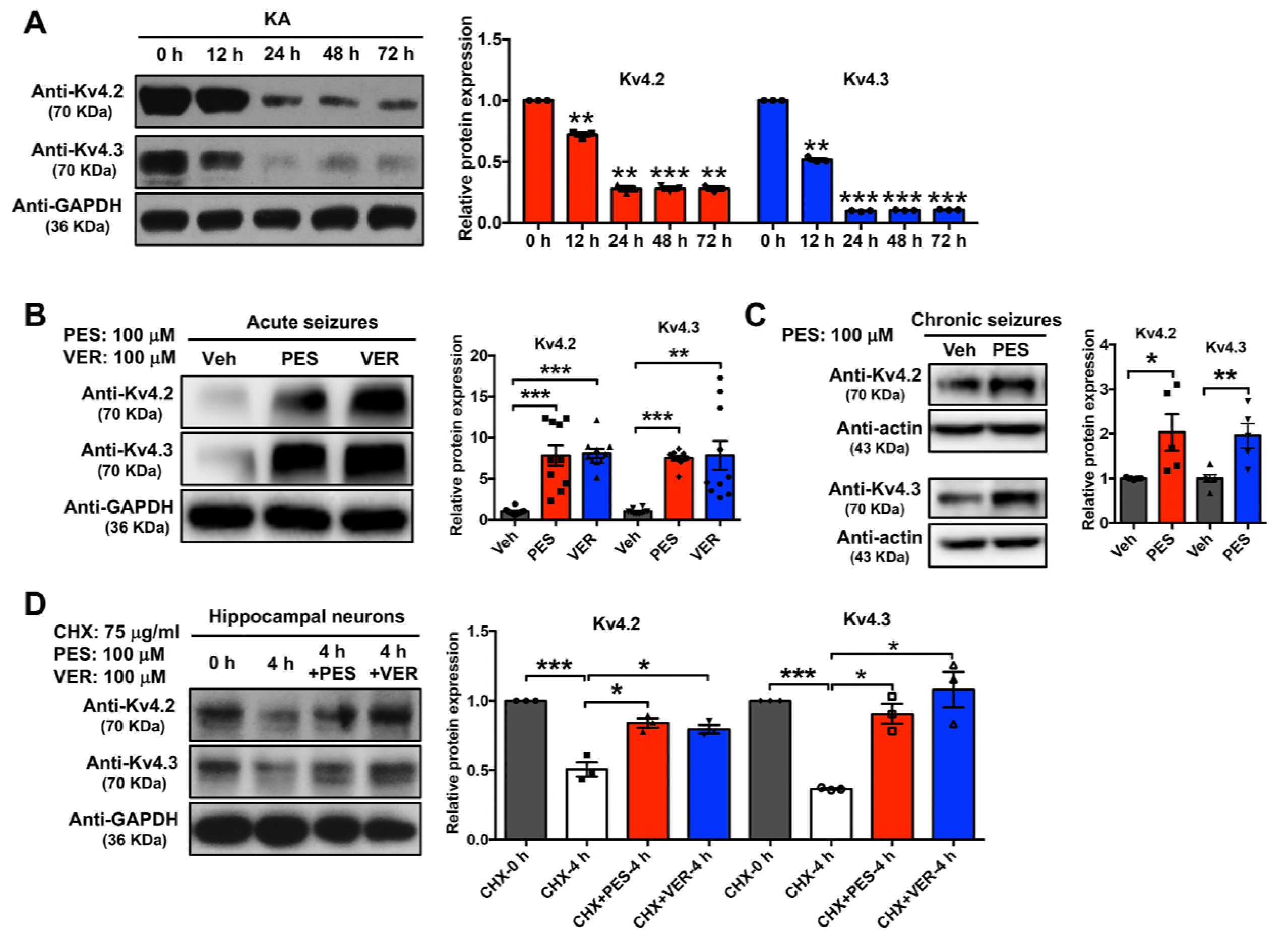
Hsp70 inhibition reduces Kv4 channel degradation caused by KA-induced seizures. **A** Western blot analysis of Kv4.2 and Kv4.3 proteins from hippocampal lysates of KA-treated rats at indicated times. Data were normalized to 0 h; n = 3 independent experiments. **B** Western blot analysis of Kv4.2 and Kv4.3 from hippocampal lysates of Veh, PES,or VER (100 μM) pre-treated rats 24 h after KA administration. Data were normalized to Veh; n = 3-10. **C** Western blot analysis of Kv4.2 and Kv4.3 from hippocampal lysates of Veh or PES (100 μM) treated rats once a day for 7 d after KA-induced chronic seizures developed. Data were normalized to Veh control; n = 5-7. **D** CHX chase assay of Kv4.2 and Kv4.3 in primary cultured hippocampal neurons. Neurons in the absence or presence of PES or VER were treated with CHX (75μg/ml), and harvested at indicated times to detect the amount of Kv4.2 and Kv4.3 in neurons by Western blotting. Western blotting of GAPDH was used as a loading control. Data were normalized to control without CHX treatment; n = 3 independent experiments. All data are expressed as the means ± s.e.m.; Comparisons for difference were analyzed using one-way ANOVA followed by Dunnet’s tests (each group compared with 0 h, Veh or CHX-4 h) for A, B and D (excluding CHX 0 h group), and unpaired Student’s t test for C and D (CHX 0 h, CHX 4 h group), \**p* <0.05; \*\**p* <0.01; \*\*\**p* <0.001.

Hsp70 and its co-chaperones have been shown to recognize and degrade wild-type and misfolded CFTR (cystic fibrosis transmembrane conductance regulator) (Meacham et al., 2001; Zhang et al., 2001) and hERG (human ether-a-go-go-related gene) channels (Li et al., 2011; Walker et al., 2010). Thus, we hypothesized that inhibition of Hsp70 increases Kv4 channel expression by preventing Kv4 from degradation. To test this hypothesis, we cultured primary rat hippocampal neurons with protein synthesis inhibitor cycloheximide (CHX, 75 μg/ml) for 4 hours in the presence or absence of PES (100 μM) or VER (100 μM), and then harvested proteins for Western blot analysis. Our results showed that both PES and VER reduced the degradation rate of either Kv4.2 or Kv4.3 α-subunits, indicating that Hsp70 facilitates the degradation of Kv4 channels in neurons (Fig. 4D).

All together, these results indicate that the degradation of Kv4 α-subunits is facilitated by upregulated Hsp70 in seizures and inhibition of Hsp70 suppresses neuronal firing by blocking Kv4 degradation caused by KA-induced seizures.

### Direct binding of Hsp70 to Kv4 auxiliary subunit KChIP4a promotes the degradation of the channel complexes

The native Kv4 channel complex is composed of Kv4 pore-forming α-subunits and auxiliary subunits including KChIPs and DPPs (Jerng & Pfaffinger, 2014). To investigate the mechanism underlying the degradation of Kv4 caused by KA-induced seizure, we performed co-immunoprecipitation (co-IP) of lysates from rat hippocampus or HEK293 cells transfected with Kv4.3-EGFP. To our surprise, Kv4 proteins were co-precipitated with Hsp70 by anti-Hsp70 antibodies from the hippocampal lysates (Fig. 5A), but not the lysates from HEK293 cells expressing Kv4.3-EGFP alone (Fig. 5B). We thus hypothesized that Hsp70 might interact with Kv4 channel complexes through its binding to auxiliary subunits, leading to the degradation of the channel complex. To test this hypothesis, we examined the interaction of Hsp70 with different KChIP members that contain a variable N-terminus and a conserved C-terminal core domain. Western blot analysis revealed that Hsp70 only interacted with KChIP4a but not KChIP1 or KChIP3 (Fig. 5C). Hsp70 was also identified to specifically interact with KChIP4a in an unbiased screening approach of co-IP combined with mass spectrometry analyses (Fig. S4A). These results suggest that the degradation of Kv4 channel complex is likely mediated by the specific binding of Hsp70 to KChIP4a containing a distinct N-terminal Kv4 inhibitory domain (KID) that causes ER retention of Kv4-KChIP4a complexes.

**Fig. 5.**
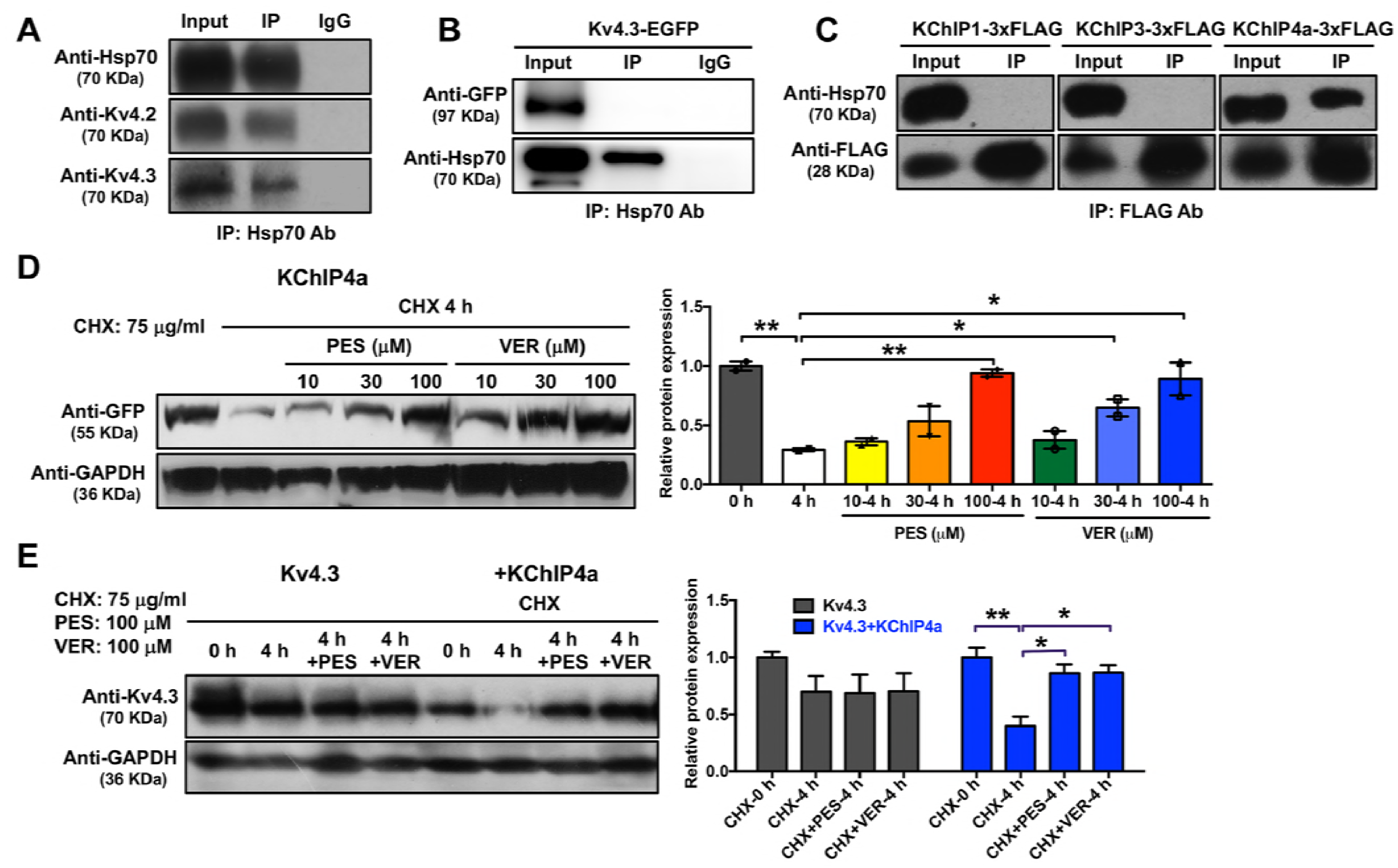
Hsp70 targets Kv4-KChIP4a channel complexes for degradation. **A** Extracted proteins from rat hippocampi were immunoprecipitated using anti-Hsp70 antibodies and analyzed by SDS-PAGE, followed by Western blot analysis for detection of Kv4.2 and Kv4.3. **B** HEK293 cells were transfected with Kv4.3-EGFP. Extracted proteins were subjected to co-IP assay using anti-Hsp70 antibodies. **C** HEK293 cells were transfected with KChIP1-3xFLAG, KChIP3-3xFLAG or KChIP4a-3xFLAG. Extracted proteins were subjected to co-IP assay using anti-Hsp70 antibodies. **D** CHX chase assay in HEK293 cells expressing KChIP4a-EGFP Cells in the absence or presence of PES or VER (10 μM, 30 μM, 100 μM) were treated with CHX (75 μg/ml), and harvested at indicated times to detect the amount of KChIP4a in cells by western blotting. Data were normalized to control without CHX treatment; n = 3 independent experiments. **E** CHX chase assay in HEK293 cells expressing Kv4.3 alone or Kv4.3 and KChIP4a-EGFP Cells were treated with CHX (75 μg/ml) in the absence or presence of PES (100 μM) or VER (100 μM), and harvested at indicated times for detection of Kv4.3 by Western blot. Data were normalized to control without CHX treatment; n = 3 independent experiments. All values are expressed as the means ± s.e.m.; comparisons for difference were analyzed using one-way ANOVA followed by Dunnet’s tests (each group compared with 4 h) for D (4 h and PES treated groups, 4 h and VER treated groups), unpaired Student’s t test for D (0 h and 4 h group), and two-way ANOVA followed by Dunnet’s tests (compared each column with CHX-4 h in each group) for E, *\*p* <0.05; *\*\*p*<0.01.

The majority of KChIP isoforms binding to Kv4 increases surface expression of Kv4 and stabilizes channel complexes (Jerng & Pfaffinger, 2014). However, the ER retention motif (N-terminal residues 12-17, LIVIVL) of KChIP4a overrides its core domain-mediated enhancement of Kv4 surface expression (Tang et al., 2013). Our results showed that the expression level of Kv4.3 co-expressed with KChIP4a was significantly lower than Kv4.3 co-expressed with KChIP1/3 (Fig. S4B), and the expression level of KChIP4a was also lower than KChIP1/3 in transfected HEK293 cells, suggesting that downregulation of Kv4 proteins by KChIP4a might result from lower stability of KChIP4a itself than KChIP1/3 (Fig. S4C). To further determine the role of KChIP4a in Hsp70 mediated degradation of Kv4-KChIP4a complexes, we evaluated protein stability of KChIP4a and Kv4.3 co-expressed with KChIP4a in the presence or absence of Hsp70 inhibitors. Both PES and VER suppressed KChIP4a-mediated degradation of KChIP4a itself and Kv4.3 proteins in a dose dependent manner (Fig. 5D-E), indicating that Hsp70 directly binds to KChIP4a and leads to the degradation of Kv4-KChIP4a channel complexes.

Given that auxiliary KChIP4a is essential for the interaction between Hsp70 and Kv4 as well as Hsp70-mediated degradation of Kv4 channels, we wondered whether KChIP4a expression was also upregulated in KA-induced seizures. Interestingly, the expression level of KChIP4a in the hippocampus was indeed increased by KA, whereas another splice variant KChIP4bl level was not altered (Fig. S4D), suggesting that the upregulation of KChIP4a is likely mediated by increased noncoding 38A-dependent alternative splicing of KChIP4, which can be triggered by inflammatory stimuli in some neuropathological conditions (Massone et al., 2011). This result also suggests that the decreased expression of Kv4 α-subunits in KA-induced seizures might result from increased expressions of Hsp70 and KChIP4a.

### The co-chaperone CHIP of Hsp70 targets KChIP4a for proteasomal degradation

Both PES and VER have been shown to suppress the activity of either constitutive Hsc70 or stress-inducible Hsp70 isoforms (Leu et al., 2009; Williamson et al., 2009). Therefore, we used the RNA interference (RNAi) assay to investigate whether these two isoforms have distinct roles in destabilizing KChIP4a. As shown in Figure 6A and 6B, both Hsp70 siRNA and Hsc70 siRNA reduced KChIP4a degradation. Knocking down the E3 ubiquitin ligase CHIP (carboxyl terminus of Hsp70-interacting protein), a co-chaperone of Hsp70/Hsc70, also inhibited KChIP4a degradation (Fig. 6B). The Hsp70/Hsc70-CHIP chaperone complex initiates the proteasomal and lysosomal degradation. Treatment with the proteasome inhibitor MG132 (15 μg/ml), but not the lysosomal inhibitor chloroquine (50 μM), resulted in inhibition of KChIP4a degradation (Fig. 6C), indicating that the Hsp70-CHIP complex directs KChIP4a toward the proteasomal degradation pathway.

**Fig. 6.**
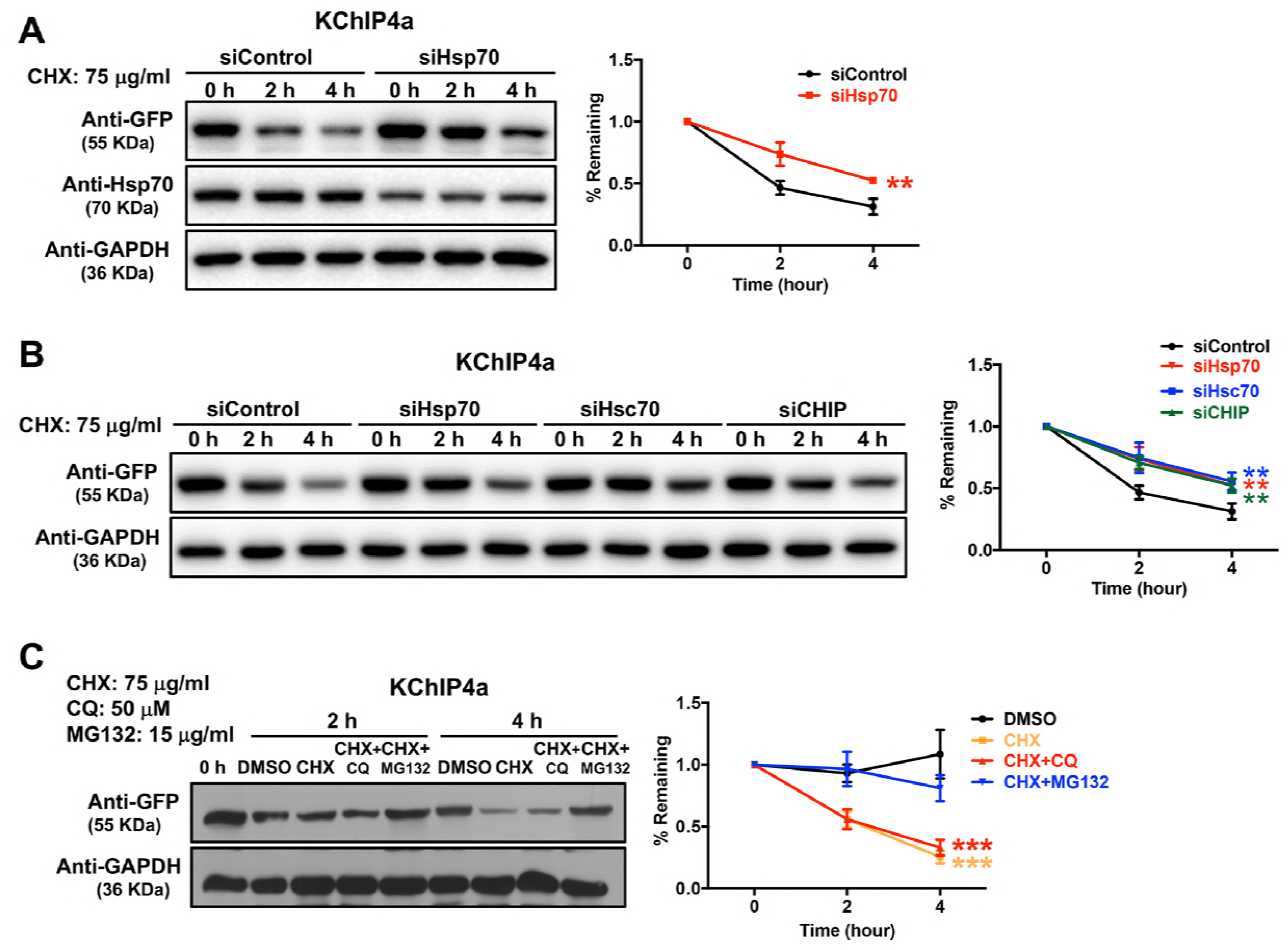
Co-chaperone CHIP of Hsp70 targets KChIP4a for proteasomal degradation. **A** CHX chase assay in HEK293 cells expressing KChIP4a-EGFP with and without silencing Hsp70. Fractions of remaining KChIP4a protein in cells incubated with control or Hsp70 siRNA plotted against time after CHX addition; n = 3 independent experiments. **B** CHX chase assay in HEK293 cells expressing KChIP4a-EGFP with and without silencing Hsp70, Hsc70 or CHIP. Fractions of remaining KChIP4a protein in cells incubated with control, Hsc70 or CHIP siRNA plotted against time after CHX addition; n = 3 independent experiments. **C** CHX chase assay in HEK293 cells expressing KChIP4a-EGFP Cells were treated with CHX (75 μg/ml) and chloroquine (CQ, 50M) or MG132 (15 μg/ml), and harvested at indicated times to detect the amount of KChIP4a in cells by Western blotting. Fractions of remaining protein plotted against time after CHX addition; n = 3 independent experiments. All values are expressed as means ± s.e.m; comparisons were analyzed using two-way ANOVA, \**\*p* <0.01; *\*\*\*p* <0.001.

### Identification of an Hsp70 recognition motif within KChIP4a for degradation

Based on the earlier data, we hypothesized that Hsp70 may directly bind to the previously identified N-terminal KID of KChIP4a and mediate the degradation of Kv4-KChIP4a complexes. To test this hypothesis, we evaluated protein stability of Kv4.3 co-expressed with KChIP4a or KChIP4aΔKID mutant in which the N-terminal KID was deleted. After 4 h of CHX (75 μg/ml) treatment, the half-life of Kv4.3 co-expressed with KChIP4a was about 2 h (Fig. 7A), whereas the half-life of Kv4.3 co-expressed with KChIP4aΔKID mutant was more than 4 h (Fig. 7A), indicating that the N-terminal KID is responsible for accelerated degradation of Kv4.3-KChIP4a complexes. Similarly, the half-life of KChIP4a was about 2 h (Fig. 7B), whereas the half-life of KChIP4aΔKID was much longer than 4 h (Fig. 7B). Next, we analyzed the effect of Hsp70 inhibitors on stability of Kv4.3 co-expressed with KChIP4aΔKID mutant in HEK293 cells. As shown in Figure 7C-D, adding PES (100 μM) or VER (100 μM) had no detectable influence on suppressing the degradation of Kv4.3 co-expressed with KChIP4aΔKID or KChIP4aΔKID mutant itself (Fig. 7C-D). Collectively, these data indicate that Hsp70 directly binds to the N-terminal KID of KChIP4a, thus leading to degradation of Kv4-KChIP4a channel complexes.

**Fig. 7.**
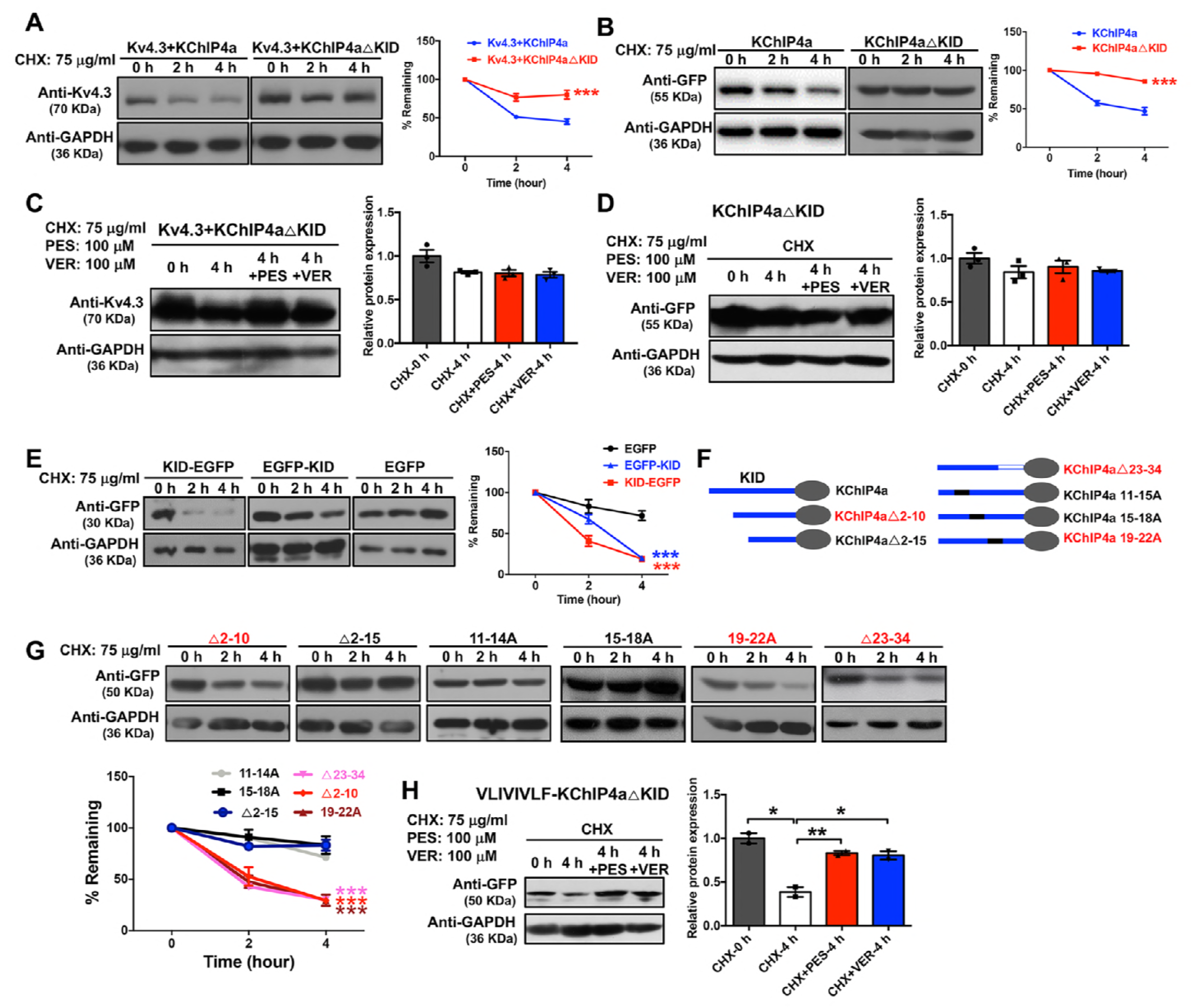
Identification of N-terminal residues VLIVIVFL of KChIP4a as a motif for degradation. **A** CHX chase assay in HEK293 cells co-transfected with Kv4.3 and KChIP4a-EGFP or Kv4.3 and KChIP4aΔKID-EGFP; n = 3 independent experiments. **B** CHX chase assay in HEK293 cells transfected with KChIP4a-EGFP or KChIP4aΔKID-EGFP; n = 3 independent experiments. **C,D** CHX chase assay in HEK293 cells expressing Kv4.3 and KChIP4aΔKID-EGFP (C) or KChIP4aΔKID-EGFP alone (D) in the absence or presence of PES (100 μM) or VER (100 μM). Data were normalized to control without CHX treatment; n = 3 independent experiments. **E** CHX chase assay in HEK293 cells transfected with KID-EGFP, EGFP-KID or EGFP; n = 3 independent experiments. **F** Schematic representation of KChIP4a mutant constructs. **G** CHX chase assay in HEK293 cells transfected with EGFP tagged KChIP4a mutants. n = 3 independent experiments. **H** CHX chase assay using HEK293 cells expressing VLIVIVFL-KChIP4aΔKID-EGFP in the absence or presence of PES (100 μM) or VER (100 μM). Data were normalized to control without CHX treatment; n =3 independent experiments. For CHX chase assay cells were treated with CHX (75 μg/ml), and harvested at indicated times to detect the amount of protein in cells by Western blotting, and fractions of remaining proteins were plotted against time after CHX addition. All data are expressed as the means ± s.e.m.; and comparisons for difference were analyzed using two-way ANOVA followed by Bonferroni’s multiple comparisons tests (compared each group with Veh) for A, B, E and G, one-way ANOVA followed by Dunnet’s tests (compared with CHX-4 h) for C, D and H (excluding CHX-0 h), and unpaired Student’s *t* test for H (CHX-0 h and CHX-4 h group), *\*p* <0.05; \**\*p* <0.01; *\*\*\*p* <0.001.

To identify the Hsp70 recognition motif within the KID of KChIP4a, we constructed various KChIP4a mutants (Fig. 7F). As shown in Figure 7E, the KID alone was sufficient to accelerate degradation when fused to either N-terminus or C-terminus of EGFP (KID-EGFP/EGFP-KID). Similar to the full-length KID, residues 11-18 (VLIVIVLF) within the KID were not only necessary but also sufficient to facilitate degradation (Fig. 7F-G), suggesting that Hsp70 targets KChIP4a for degradation through binding to the VLIVIVLF motif. Treatment with Hsp70 inhibitor PES (100 μM) or VER (100 μM) and the proteasome inhibitor MG132, but not the lysosomal inhibitor chloroquine, reduced the VLIVIVLF motif-mediated degradation (Fig. 7H and S5). These data confirm the residues VLIVIVLF within KChIP4a functioning as a motif for recognition of Hsp70 and Hsp70-CHIP assisted degradation of Kv4-KChIP4a complexes.

## DISCUSSION

The goal of this study is to investigate whether Hsp70 is involved in pathogenesis of epilepsy. Although it is well known that Hsp70 is upregulated as a stress biomarker in various types of epilepsy (Kandratavicius et al., 2014; Lively & Brown, 2011; Rejdak et al., 2012; Yang et al., 2008), the role of Hsp70 in epileptogenesis is still unclear and controversial. This study reveals an unexpected deleterious role of Hsp70 in seizures, and also provides evidence for pharmacological intervention of seizures through restoring expression and function of Kv4 channels by inhibiting endogenous Hsp70. Our findings indicate that KA induced overexpression of Hsp70 exacerbates seizure pathology and increases neuronal hyperactivity by promoting degradation of Kv4 channel complexes. This Hsp70-facilitated degradation of Kv4 channel complexes is dependent on the presence of inhibitory auxiliary subunit KChIP4a that directly interacts with Hsp70 through its unique N-terminal Kv4 channel inhibitory domain (KID) (Fig. 8).

**Fig. 8.**
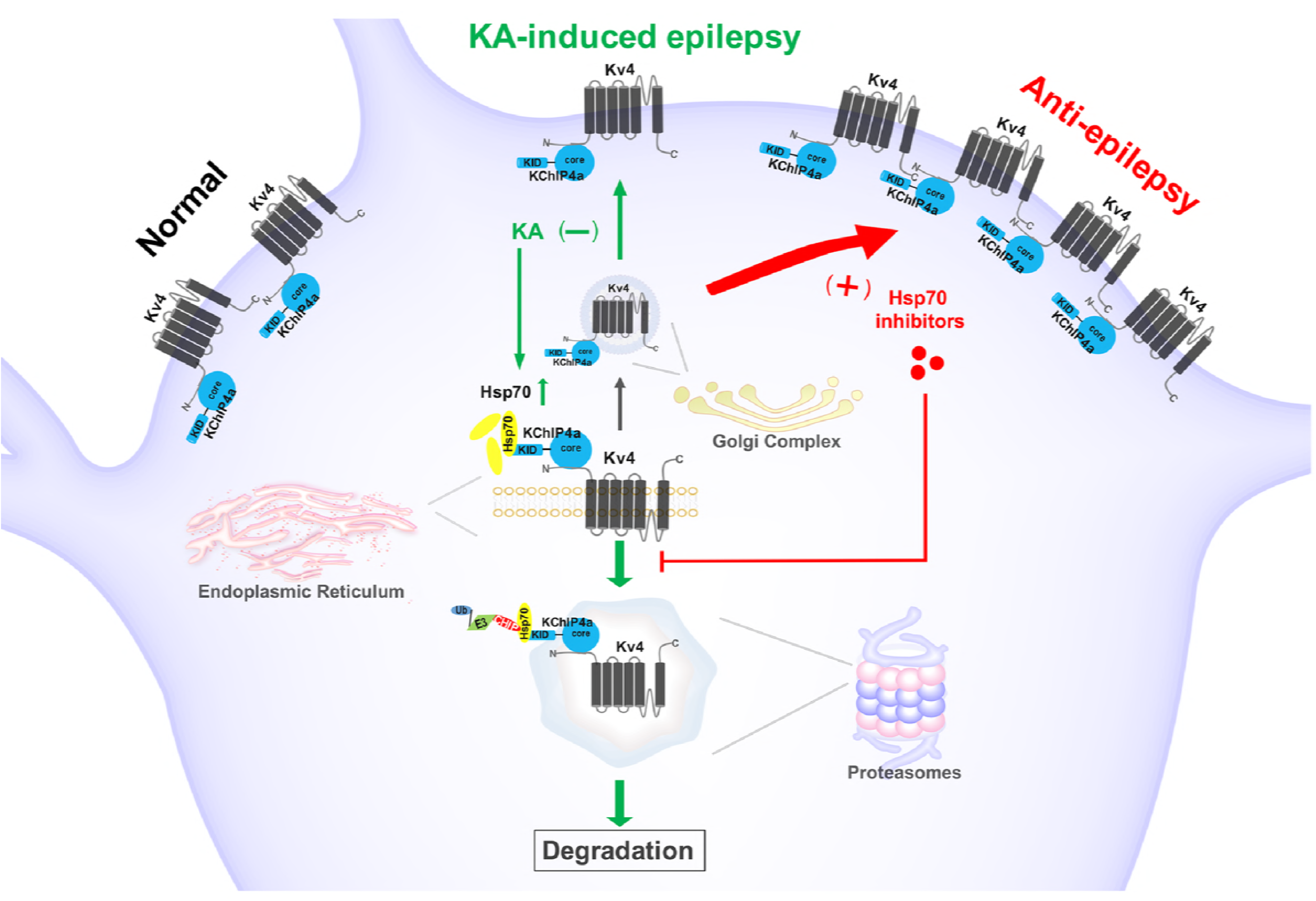
The proposed model for the role of Hsp70 in regulation of neuronal excitability and epilepsy. KChIP4a binds to the N-terminus of Kv4 forming the Kv4-KChIP4a channel complex. Hsp70/Hsc70 binds to the N terminal KID of KChIP4a, and promotes proteasomal degradation of Kv4-KChIP4a complexes dependent on the co-chaperone/E3 ubiquitin ligase CHIP. KA stimulates Hsp70/Hsc70 expression, which promotes the degradation of Kv4-KChIP4a complexes and leads to reduced surface expression of Kv4 channels and decreased *I_SA_* current, resulting in neuronal hyperexcitability or epilepsy (pathway shown in green). Hsp70 inhibition suppresses degradation of Kv4-KChIP4a complexes mediated by upregulated Hsp70, which maintains surface expression of Kv4 channels for normal neuronal activity (Anti-epilepsy pathway shown in red). KA: kainic acid; KID: Kv4 channel inhibitory domain; core: KChIP-conserved C-terminal domain; CHIP: carboxyl terminus of Hsp70-interacting protein; Ub: ubiquitin; E3: E3 ubiquitin ligase; N: N-terminus; C: C-terminus.

Kv4 channels underlie the main somatodendritic A-type Kv currents in most hippocampal pyramidal neurons and play a critical role in regulating suprathreshold dendritic signals such as dendritic action potential backpropagation, furthering dendritic Ca^2+^ influx and excitability (Chen et al., 2006). The relevance of Kv4.2 to epilepsy is supported by the identification of a C-terminal truncation mutation of Kv4.2 in a patient with TLE (Singh et al., 2006), and the observations from Kv4.2 knockout mice showing increased susceptibility to seizures induced by KA (Barnwell et al., 2009). Studies in animal epilepsy models also demonstrate a reduction of Kv4.2 mRNA in the dentate granule cells of the hippocampus (Tsaur et al., 1992), and an activity-dependent reduction of Kv4 A-type currents (Bernard et al., 2004), which might result from increased Kv4.2 internalization (Kim et al., 2007), and microRNA miR-324-5p mediated downregulation of Kv4.2 (Gross et al., 2016). Here we present a novel molecular mechanism by which Hsp70-CHIP regulates Kv4 degradation in response to acute and chronic seizures.

We previously demonstrated that KChIP4a is capable of retaining not only itself but also interacting with Kv4 α-subunits in the endoplasmic reticulum (ER) through its N-terminal KID, leading to subsequent clearance of Kv4 channels by the ER-associated degradation (ERAD) system (Tang et al., 2013). In this study, our results provide mechanistic insights into the cytosolic chaperone pathways that regulate the expression of Kv4-KChIP4a channel complexes. Our data show that auxiliary KChIP4a subunit controls a sorting step in the endoplasmic reticulum that diverts channels from anterograde transport to the ERAD complex (Fig. 8). This likely involves multiple KChIP4a-dependent steps that include the association of Kv4 α-subunit with the KChIP-conserved C-terminal core domain, the interaction of the Hsp70-CHIP chaperone complex with the N-terminal KID of KChIP4a, and perhaps ubiquitination of KChIP4a and/or Kv4 subunits by CHIP. Notably, ubiquitination may not be necessary for proteasomal degradation of Kv4 α-subunits. In our study, it is likely that KChIP4a, ubiquitinated by co-chaperone/E3 ubiquitin ligase CHIP, acts as an adaptor that targets Kv4 ?-subunit for degradation. Similar to Hsp70, the expression of KChIP4a, but not Hsp70-binding-defective KChIP4 splicing variant KChIP4bl, is also induced in KA-induced seizures, which results in the increase of Kv4-KChIP4a complex formation, thus leading to higher likelihood of ubiquitination and degradation of Kv4 channels. The seizure activity dependent upregulation of KChIP4a is probably driven by RNA polymerase III-dependent noncoding RNA 38A that has been reported in Alzheimer’s disease for driving shift of alternative splicing of KChIP4 mRNAs under inflammation, and promoting the synthesis of KChIP4a isoform (Massone et al., 2011).

We previously reported that different KChIPs, including KID-containing KChIPs and KChIPs without KID, could compete for heteromultimeric assembly with Kv4 α-subunits, which depends on the relative expression level of each KChIP subunit (Zhou et al., 2015). Therefore, although the stoichiometry of the KChIP4a-Hsp70 complex is unclear, a Kv4-KChIPs complex containing even only one KChIP4a molecule may still be recognized by Hsp70-CHIP for degradation. Moreover, besides KChIP4a, two previously identified brain KChIP isoforms, KChIP2x and KChIP3x, also contain the VLIVIVLF like Hsp70 recognition motif and may promote the degradation of Kv4 channels (Fig. S6), further increasing the likelihood of Kv4-KChIPs degradation. Given that inhibiting Hsp70 may increase the risk of neuronal death (Chang et al., 2014), developing small-molecule compounds aimed at specifically disrupting the interaction between KChIP4a and Hsp70 may provide a beneficial potential for epilepsy.

Although neuroprotective roles of Hsp70 have been implicated in various neurodegenerative diseases such as the polyQ diseases (Adachi et al., 2003; Kazemi-Esfarjani & Benzer, 2000; Wang et al., 2013), ALS (Bruening et al., 1999; Kieran et al., 2004) and PD (Auluck et al., 2002; Klucken et al., 2004), activation of Hsp70 is not beneficial in all instances. For example, pharmacological up-regulation of Hsp70 by celastrol induces neuronal death in a primary cellular model of motoneuron neurodegeneration (Kalmar & Greensmith, 2009), and Hsp70 is associated with neurodegeneration and immunological deregulation in multiple sclerosis (MS) through promoting or exacerbating an immunological response mediated by its cytokine-like property as well as its myelin-peptide adjuvant capacity (Asea et al., 2000; Asea et al., 2002; Mycko et al., 2008). At the present, it is unclear if roles of Hsp70 in epilepsy are beneficial or harmful. Here we show that seizure-induced upregulation of Hsp70 is likely to play a pathological role in enhancement of neuronal hyperexcitability.

Both constitutive (Hsc70) and stress-inducible (Hsp70) isoforms of cytosolic Hsp70 subfamily members have been proposed to regulate the intracellular trafficking and quality control of ion channels (Young, 2014). Goldfarb et al show that moderate overexpression of Hsp70 increases the expression of the epithelial sodium channel (ENaC) by promoting protein folding, whereas overexpressing large amounts of either Hsc70 or Hsp70 decreases ENaC by increasing its ubiquitination and degradation (Goldfarb et al., 2006). In this study, our results also suggest that constitutive Hsc70 or induced Hsp70 can play a similar role in facilitating degradation of Kv4-KChIP4a complexes in KA-induced seizures, which is dependent on the co-chaperone/E3 ubiquitin ligase CHIP. Two small molecule inhibitors of Hsp70 used in this study, PES and VER, have been shown to dually target to both Hsp70 and Hsc70 proteins(Schlecht et al., 2013; Williamson et al., 2009). The PES compound binds to the substrate binding domain (SBD) of Hsp70/Hsc70 and reversibly disrupts the association of Hsp70 with Hsp90 and co-chaperones (Kalmar & Greensmith, 2009; Leu et al., 2009). The ATP analog VER targets to the nucleotide binding domain (NBD) of Hsp70/Hsc70 and thereby acts as an ATP-competitive inhibitor preventing allosteric control between NBD and SBD (Schlecht et al., 2013). Recently, another small molecule inhibitor of ATPase activity of Hsp70/Hsc70, apoptozole (Az), has been proposed to suppress ubiquitination of ΔF508-CFTR mutant by disrupting interaction of the mutant with Hsc70 and CHIP, and, as a consequence, to promote cell surface trafficking of the mutant (Cho et al., 2011). All of these observations are consistent with our findings that are indicative of a similar underlying mechanism by which Hsp70/Hsc70 inhibitors can ameliorate seizure severity.

In summary, we show a previously unrecognized but important role of Hsp70 in KA-induced seizures. Our findings suggest that an alternative therapeutic strategy through inhibition of Hsp70/Hsc70 chaperone systems and enhancement of Kv4 function may be beneficial for treatment of epilepsy.

## MATERIALS AND METHODS

### Plasmid Construction

For Western blotting analysis, the cDNAs of human Kv4.3 (NG_032011.2), human KChIP1 (NP_075218), rat KChIP3 (NM_032462.2), mouse KChIP4a (NP_084541) and KChIP4a mutants were cloned into a pEGFP-N1 or pEGFP-C2 vector. For co-immunoprecipitation analysis, the cDNAs of KChIP1, KChIP3 and KChIP4a were inserted into the vector pcDNA3.1(+) with three tandem-repeated FLAG epitopes (DYKDDDDK) at the C-terminus.

### Isolation and primary culture of rat hippocampal neurons

Hippocampal explants isolated from embryonic day 18 rats of either sex were digested with 0.25% trypsin for 30 min at 37 °C, followed by triturating with a pipette in plating medium (DMEM with 10% FBS). Dissociated neurons were plated onto 35 mm dishes coated with poly-D-lysine (0.5 mg/ml, Sigma Aldrich) at a density of 5 x 10^5^ cells per dish. After 4 h, the medium was changed to Neurobasal medium supplemented with 2% B27 and 0.5 mM GlutaMAX-I (Life Technologies). Hippocampal neurons were cultured for 5-7 days before electrophysiological recordings or Western blotting assay.

### Animals

Adult male Sprague-Dawley rats (190-210 g) used in this study are from the Department of Experimental Animal Sciences, Peking University Health Science Center. All animals were allowed free access to food and water. The experimental protocols were approved by the Animal Use and Care Committee of Qingdao University.

### Surgery and intracerebroventricular (ICV) administration of Hsp70 inhibitors

Rats were anaesthetized with 400 mg/kg of chloral hydrate i.p. and placed on stereotaxic apparatus. Three cortical skull-mounted electroencephalography (EEG) electrodes were attached. For intracerebroventricular (ICV) microinjections, the stainless-steel guide cannula (RWD Life Science) was implanted into the lateral ventricle of the brain. The coordinates were as follows: AP = 0.8 mm posterior to bregma, ML = 1.5 mm lateral to bregma, DV = 3.8 mm below the surface of the skull. EEG electrodes and guide cannula were then secured with dental cement (Jimenez-Pacheco et al., 2013). 7 days after surgery, rats were gently restrained and an infusion cannula (RWD Life Science) was inserted into the lateral ventricle through the guide cannula to a depth of 3.8 mm below the surface of the skull. 2-phenylethynesulfonamide (PES, Sigma Aldrich, P0122) and VER-155008 (VER, Sigma Aldrich, SML0271) were dissolved in dimethyl sulfoxide (DMSO, Sigma Aldrich) at 50 mM and stored at −20 °C until use. PES and VER (10, 30, or 100 μM) were diluted in 1 μl ACSF (125 mM NaCl, 2.5 mM KCl, 21.4 mM NaHCO_3_, 1.25 mM NaH_2_PO_4_, 2.0 mM CaCl_2_, 1.0 mM MgCl_2_, and 11.1 mM glucose) and was infused into the lateral ventricle in 4 min (0.25 μl/min) followed by an additional 1 min to allow diffusion before the injection needle was removed. Rats in vehicle control were infused with 1μl ACSF.

### Spontaneous locomotor activity test

To evaluate spontaneous locomotor activity, we used an activity monitor (Experimental Factory of Chinese Academy of Medical Sciences, Beijing, China) consisting of 4 plexiglass cylinders each equipped with three infrared beams and an automated counting system. Briefly, animals were individually placed in the cylinder for 3 min for adaptation to the new environmental. Locomotor activity was then assessed by counting the number of infrared beam crossings in the photocell apparatus in 5 min.

### Acute model of seizures and EEG recordings

7 days after surgery, PES and VER were administrated ICV for 7 days (once a day) before KA injection. In the vehicle group, rats injected with DMSO in ACSF under the same regimen were used as controls. 24 h after the last administration, 1 μg kainic acid (KA) (dissolved in 1 μl ACSF) was injected into the lateral ventricle of the brain. Subsequently, rats were individually housed and recorded for 1 h by electroencephalogram (EEG) telemetry under video monitoring. EEG telemetry (BIOPAC Systems) and video monitoring (Hikvision Digital Technology) were started 15 min before KA injection for baseline EEG recordings and behavioral activity. 1 h after KA injection, seizures were terminated at 1 h after onset with the use of sodium pentobarbital (SP; 30 mg/kg, s.c.; Sigma-Aldrich) (Huang et al., 2012) if necessary. All procedures concerning animals were approved by the Animal Use and Care Committee of Qingdao University.

For quantitative analysis of EEG recordings, EEG data were uploaded into AcqKnowledge 4.2.1 software (BIOPAC Systems) for power of whole bandwidth analysis by using “EEG Frequency Analysis” function. Coastline index was determined as the cumulative difference between successive points on the EEG (White et al., 2006). OriginPro version 8.6 (OriginLab) was used for extracted coastline index.

For seizure behavioral scoring, the epileptic behaviors from video recordings were scored using the following Racine’s scale (Racine, 1972): stage 1, facial myoclonus; stage 2, facial and mild forelimb clonus; stage 3, severe forelimb clonus; stage 4, rearing in addition to severe forelimb clonus; stage 5, rearing and falling in addition to severe forelimb clonus.

### Chronic model of seizures and Video & EEG recordings

After unilateral intracerebroventricular (ICV) injection (AP = 0.8 mm posterior to bregma, ML = 1.5 mm lateral to bregma, DV = 3.8 mm below the surface of the skull) of 1 μg KA, rats experienced at least 2 h of convulsive SE, and then chronic seizures developed within 2-3 weeks. After chronic seizures developed, each rat was implanted with a guide cannula for ICV administration and a sender (Chengdu Technology & Market Corp., LTD) for EEG recording. After a 3-4 d recovery period, the baseline of spontaneous seizures was recorded for 7-10 consecutive days. The selected rats were then ICV administrated with PES (100 μM) or DMSO (vehicle control) once a day and monitored continuously for 7 d for SRSs by video and EEG recording. For EEG recording, animals were housed individually cages, each fitted with a receiver and EEG was recorded and analyzed using BW-200 Data acquisition system for life science by wireless (Chengdu Technology & Market Corp., LTD). Seizure frequency was measured according to video recording and identified by two trained experimenters blinded to the experimental conditions.

### Co-immunoprecipitation

For Co-immunoprecipitation (Co-IP) assays, transfected HEK 293 cells were washed three times with ice cold phosphate buffered saline. Rat hippocampus was prepared by sonicating samples for 10 s. Cells and prepared hippocampus were lysed with lysis buffer (150 mM NaCl, 20 mM Tris, 1% Triton X-100, 1% sodium deoxycholate, 10 mM EDTA, and proteinase inhibitor mixture (Roche Applied Science), pH 8.0) at 4 °C for 30 min. The lysates were then centrifuged at 13,000 × g for 10 min to yield the protein extract in the supernatant. One fraction containing 200 mg of protein was incubated with 1 ~ 2 ml ANTI-FLAG M2 Affinity Gel (Sigma) or 50 μl of Protein A/G PLUS-Agarose (Santa Cruz Biotech) and 1 mg Hsp70 antibody (Abcam ab2787) for 5 h or overnight at 4 °C to immunoprecipitate proteins, and the other fraction was prepared as input protein. After incubation, co-IP were extensively washed four times with Tris buffered saline and eluted with 0.1 M glycine HCI, pH 3.5. The input and IP proteins were both subjected to Western blotting analysis.

### Mass spectrometry analysis for protein interactions

HEK 293 cells were transfected with KChIP4a-FLAG, KChIP4a-core-FLAG, KChIP3-FLAG or control plasmid. After 24 hours of transfection, cells were lysed with lysis buffer (150 mM NaCl, 20 mM Tris, 1% Triton X-100, 1% sodium deoxycholate, 10 mM EDTA, and proteinase inhibitor mixture (Roche Applied Science), pH 8.0) at 4 °C for 30 min. The lysates were then centrifuged at 13,000 × g for 10 min to yield the protein extract in the supernatant. All the supernatants were incubated with ANTI-FLAG M2 Affinity Gel (Sigma) overnight at 4 °C. After incubation, the agarose beads were washed for four times with Tris buffered saline and proteins were eluted with 0.1 M glycine HCI, pH 3.0. The eluted proteins were loaded on SDS-PAGE before stained with Coomassie blue for 1-2 hours and de-stained with de-stain buffer for three times. For Mass spectrometry analysis, the gel was placed on a light box and the protein bands non-existent in KChIP3-FLAG-IP lane were excised. Mass spectrometry analysis was performed at the Teaching Center of Biology Experiment, School of Life Sciences, Sun Yat-Sen University (Guangzhou, China).

### RNA interference

All siRNA were synthesized by Guangzhou Ribobio co., LTD. The sequences were as follows: human Hsp70 sense: CGGACAAGAAGAAGGUGCUdTdT; antisense: dTdTGCCUGUUCUUCUUCCACGA. human Hsc70 sense:GCAACTGTTGAAGATGAGAAAdTdT; antisense: dTdCGUUGACAACUUCUACUCUUU. human CHIP sense: GCAGTCTGTGAAGGCGCACTTdTdT; antisense: dTdCGUCAGACACUUCCGCGUGAA. HEK293 cells were transfected with siRNA 24 ~ 48 h before transfected with plasmids.

### Cycloheximide Treatment

For cycloheximide (CHX) treatment, transfected HEK293 cells were treated with CHX for various time periods (0 h, 2 h, 4 h) before cells were lysed with lysis buffer (150 mM NaCl, 20 mM Tris, 1% Triton X-100, 1% sodium deoxycholate, 0.1% SDS, 10 mM EDTA, and proteinase inhibitor mixture (Roche Applied Science), pH 8.0) at 4 °C for 30 min. The cell lysates were then centrifuged at 13,000 × g for 10 min to yield protein extracts in the supernatant. The supernatants were then subjected to Western blotting analysis.

### Western Blotting Assay

For Western blotting assay, protein samples were loaded on SDS-PAGE and transferred onto PVDF membranes (Millipore). After blocking, membranes were incubated with rabbit monoclonal anti-Hsp70 (1:2000; Abcam, ab2787), rabbit monoclonal anti-GFP (1:2000; Abcam, ab1218), mouse monoclonal anti-FLAG (1:4000; Sigma, F3165), mouse monoclonal anti-Kv4.2 (1:2000; Abcam, ab99040), mouse monoclonal anti-Kv4.3 (1:2000; Abcam, ab99045) or mouse monoclonal anti-GAPDH (1:10000; Santa Cruz, sc-32233) at 4 °C overnight. The membranes were then incubated with their corresponding secondary HRP-conjugated antibodies and detected using an ECL Western blotting detection system (Millipore). Detection of signals was calculated using Quantity One software (Bio-Rad). For quantification, the signals from anti-Hsp70, anti-Kv4.2, anti-Kv4.3 or anti-GFP over anti-GAPDH were normalized.

### Electrophysiological Recordings

For whole-cell patch clamp recordings in neurons, currents were recorded at room temperature using the EPC 10 USB amplifier with PatchMaster software (HEKA Electronics). Patch pipettes were pulled from borosilicate glass and fire-polished to a resistance of 2 ~ 4 megaohms. The bath solution contained 125 mM NaCl, 2.5 mM KCl, 25 mM NaHCO_3_, 2 mM CaCl_2_, 2 mM MgCl_2_, at pH 7.4 (for isolating *I_SA_*, with 1 μM tetrodotoxin). The pipette solution contained 120 mM K-gluconate, 20 mM KCl, 5 mM NaCl, 10 mM HEPES, 4 mM Mg-ATP, 3 mM Tris-GTP, 14 mM phosphocreatine, at pH 7.4. Cells were held at −100 mV for *I*_*SA*_ recording and voltage steps (ranging from −70 mV to 50 mV, 10 mV/step) from −100 mV and −30 mV (prepulse) with duration of 1000 ms were applied to activate whole-cell currents. Subtraction of the −30 mV traces from the −100 mV traces revealed *I_SA_* current. For current-clamp recordings, cells were held at 0 pA, and the firing rates of neurons were measured by a series of 1000-ms depolarizing current injection in 5-pA steps from 0 pA to 45 pA. The first action potential of a train was used to determine the threshold, defined as the voltage at which the first derivative of the membrane potential increased by 10 V/s. Data were acquired using PatchMaster software (HEKA Electronics). OriginPro version 8.6 (OriginLab) was used for data analysis.

### Data Analysis and Statistics

All data are presented as mean ± SEM. Comparisons between two groups were analyzed using unpaired Student’s *t*-tests, while multi-group comparisons were analyzed using two-way ANOVA followed by Bonferroni’s multiple comparisons tests or one-way ANOVA followed by Dunnet’s tests. Data marked with asterisks are significantly different from the control with p-values as follows: \**p*< 0.05, \*\**p*< 0.01, \*\*\**p*< 0.001.

## AUTHOR CONTRIBUTIONS

F.H, J.H.Z., Y.Q.T. and K.W.W. conceived and designed experiments; F.H, J.H.Z., Y.X.L., L.Z.G, Z.H. and Y.Q.T performed experiments and analyzed the data; N.N.W assisted in electrophysiological recordings. F.H., J.H.Z., Y.Q.T. and K.W.W. wrote the manuscript.

## ACKNOWLEDGMENTS

We would like to thank C. Qi, T. Ma and G. Wang for technical assistance and M. Li for discussion. K.W.W. wishes to thank J. M. Wang for her consistent support during this research. This work was supported by research grants to K.W.W. from the National Science Foundation of China (31370741 and 81573410), and the Ministry of Science and Technology of China (2013CB531302).

## REFERENCES

Abisambra J, Jinwal UK, Miyata Y, Rogers J, Blair L, Li X, Seguin SP, Wang L, Jin Y, Bacon J, Brady S, Cockman M, Guidi C, Zhang J, Koren J, Young ZT, Atkins CA, Zhang B, Lawson LY, Weeber EJ et al. (2013) Allosteric heat shock protein 70 inhibitors rapidly rescue synaptic plasticity deficits by reducing aberrant tau. Biol Psychiatry 74: 367-74

Adachi H, Katsuno M, Minamiyama M, Sang C, Pagoulatos G, Angelidis C, Kusakabe M, Yoshiki A, Kobayashi Y, Doyu M, Sobue G (2003) Heat shock protein 70 chaperone overexpression ameliorates phenotypes of the spinal and bulbar muscular atrophy transgenic mouse model by reducing nuclear-localized mutant androgen receptor protein. J Neurosci 23: 2203-11

An WF, Bowlby MR, Betty M, Cao J, Ling HP, Mendoza G, Hinson JW, Mattsson KI, Strassle BW, Trimmer JS, Rhodes KJ (2000) Modulation of A-type potassium channels by a family of calcium sensors. Nature 403: 553-6

Asea A, Kraeft SK, Kurt-Jones EA, Stevenson MA, Chen LB, Finberg RW, Koo GC, Calderwood SK (2000) HSP70 stimulates cytokine production through a CD14-dependant pathway, demonstrating its dual role as a chaperone and cytokine. Nat Med 6: 435-42

Asea A, Rehli M, Kabingu E, Boch JA, Bare O, Auron PE, Stevenson MA, Calderwood SK (2002) Novel signal transduction pathway utilized by extracellular HSP70: role of toll-like receptor (TLR) 2 and TLR4. J Biol Chem 277: 15028-34

Auluck PK, Chan HY, Trojanowski JQ, Lee VM, Bonini NM (2002) Chaperone suppression of alpha-synuclein toxicity in a Drosophila model for Parkinson’s disease. Science 295: 865-8

Barnwell LF, Lugo JN, Lee WL, Willis SE, Gertz SJ, Hrachovy RA, Anderson AE (2009) Kv4.2 knockout mice demonstrate increased susceptibility to convulsant stimulation. Epilepsia 50: 1741-51

Bernard C, Anderson A, Becker A, Poolos NP, Beck H, Johnston D (2004) Acquired dendritic channelopathy in temporal lobe epilepsy. Science 305: 532-5

Bruening W, Roy J, Giasson B, Figlewicz DA, Mushynski WE, Durham HD (1999) Up-regulation of protein chaperones preserves viability of cells expressing toxic Cu/Zn-superoxide dismutase mutants associated with amyotrophic lateral sclerosis. J Neurochem 72: 693-9

Burgoyne RD (2007) Neuronal calcium sensor proteins: generating diversity in neuronal Ca2+ signalling. Nat Rev Neurosci 8: 182-93

Chang BS, Lowenstein DH (2003) Epilepsy. N Engl J Med 349: 1257-66

Chang CC, Chen SD, Lin TK, Chang WN, Liou CW, Chang AY, Chan SH, Chuang YC (2014) Heat shock protein 70 protects against seizure-induced neuronal cell death in the hippocampus following experimental status epilepticus via inhibition of nuclear factor-kappaB activation-induced nitric oxide synthase II expression. Neurobiol Dis 62: 241-9

Chen X, Yuan LL, Zhao C, Birnbaum SG, Frick A, Jung WE, Schwarz TL, Sweatt JD, Johnston D (2006) Deletion of Kv4.2 gene eliminates dendritic A-type K+ current and enhances induction of long-term potentiation in hippocampal CA1 pyramidal neurons. J Neurosci 26: 12143-51

Cho HJ, Gee HY, Baek KH, Ko SK, Park JM, Lee H, Kim ND, Lee MG, Shin I (2011) A small molecule that binds to an ATPase domain of Hsc70 promotes membrane trafficking of mutant cystic fibrosis transmembrane conductance regulator. J Am Chem Soc 133: 20267-76

Ekimova IV, Nitsinskaya LE, Romanova IV, Pastukhov YF, Margulis BA, Guzhova IV (2010) Exogenous protein Hsp70/Hsc70 can penetrate into brain structures and attenuate the severity of chemically-induced seizures. J Neurochem 115: 1035-44

Goldfarb SB, Kashlan OB, Watkins JN, Suaud L, Yan W, Kleyman TR, Rubenstein RC (2006) Differential effects of Hsc70 and Hsp70 on the intracellular trafficking and functional expression of epithelial sodium channels. Proc Natl Acad Sci U S A 103: 5817-22

Gross C, Yao X, Engel T, Tiwari D, Xing L, Rowley S, Danielson SW, Thomas KT, Jimenez-Mateos EM, Schroeder LM, Pun RYK, Danzer SC, Henshall DC, Bassell GJ (2016) MicroRNA-Mediated Downregulation of the Potassium Channel Kv4.2 Contributes to Seizure Onset. Cell Rep 17: 37-45

Hartl FU, Bracher A, Hayer-Hartl M (2011) Molecular chaperones in protein folding and proteostasis. Nature 475: 324-32

Huang Z, Lujan R, Martinez-Hernandez J, Lewis AS, Chetkovich DM, Shah MM (2012) TRIP8b-independent trafficking and plasticity of adult cortical presynaptic HCN1 channels. J Neurosci 32: 14835-48

Jerng HH, Pfaffinger PJ (2014) Modulatory mechanisms and multiple functions of somatodendritic A-type K (+) channel auxiliary subunits. Front Cell Neurosci 8: 82

Jimenez-Pacheco A, Mesuret G, Sanz-Rodriguez A, Tanaka K, Mooney C, Conroy R, Miras-Portugal MT, Diaz-Hernandez M, Henshall DC, Engel T (2013) Increased neocortical expression of the P2X7 receptor after status epilepticus and anticonvulsant effect of P2X7 receptor antagonist A-438079. Epilepsia 54: 1551-61

Jinwal UK, Miyata Y, Koren J, 3rd, Jones JR, Trotter JH, Chang L, O’Leary J, Morgan D, Lee DC, Shults CL, Rousaki A, Weeber EJ, Zuiderweg ER, Gestwicki JE, Dickey CA (2009) Chemical manipulation of hsp70 ATPase activity regulates tau stability. J Neurosci 29: 12079-88

Kalmar B, Greensmith L (2009) Activation of the heat shock response in a primary cellular model of motoneuron neurodegeneration-evidence for neuroprotective and neurotoxic effects. Cell Mol Biol Lett 14: 319-35

Kandratavicius L, Hallak JE, Carlotti CG, Jr., Assirati JA, Jr., Leite JP (2014) Hippocampal expression of heat shock proteins in mesial temporal lobe epilepsy with psychiatric comorbidities and their relation to seizure outcome. Epilepsia 55: 1834-43

Kazemi-Esfarjani P, Benzer S (2000) Genetic suppression of polyglutamine toxicity in Drosophila. Science 287: 1837-40

Kieran D, Kalmar B, Dick JR, Riddoch-Contreras J, Burnstock G, Greensmith L (2004) Treatment with arimoclomol, a coinducer of heat shock proteins, delays disease progression in ALS mice. Nat Med 10: 402-5

Kim J, Jung SC, Clemens AM, Petralia RS, Hoffman DA (2007) Regulation of dendritic excitability by activity-dependent trafficking of the A-type K+ channel subunit Kv4.2 in hippocampal neurons. Neuron 54: 933-47

Kim J, Wei DS, Hoffman DA (2005) Kv4 potassium channel subunits control action potential repolarization and frequency-dependent broadening in rat hippocampal CA1 pyramidal neurones. J Physiol 569: 41-57

Klucken J, Shin Y, Masliah E, Hyman BT, McLean PJ (2004) Hsp70 Reduces alpha-Synuclein Aggregation and Toxicity. J Biol Chem 279: 25497-502

Leu JI, Pimkina J, Frank A, Murphy ME, George DL (2009) A small molecule inhibitor of inducible heat shock protein 70. Mol Cell 36: 15-27

Li P, Ninomiya H, Kurata Y, Kato M, Miake J, Yamamoto Y, Igawa O, Nakai A, Higaki K, Toyoda F, Wu J, Horie M, Matsuura H, Yoshida A, Shirayoshi Y, Hiraoka M, Hisatome I (2011) Reciprocal control of hERG stability by Hsp70 and Hsc70 with implication for restoration of LQT2 mutant stability. Circ Res 108: 458-68

Lively S, Brown IR (2011) Induction of heat shock proteins in the adult rat cerebral cortex following pilocarpine-induced status epilepticus. Brain Res 1368: 271-80

Loscher W, Klitgaard H, Twyman RE, Schmidt D (2013) New avenues for anti-epileptic drug discovery and development. Nat Rev Drug Discov 12: 757-76

Luders J, Demand J, Hohfeld J (2000) The ubiquitin-related BAG-1 provides a link between the molecular chaperones Hsc70/Hsp70 and the proteasome. J Biol Chem 275: 4613-7

Lugo JN, Barnwell LF, Ren Y, Lee WL, Johnston LD, Kim R, Hrachovy RA, Sweatt JD, Anderson AE (2008) Altered phosphorylation and localization of the A-type channel, Kv4.2 in status epilepticus. J Neurochem 106: 1929-40

Massone S, Vassallo I, Castelnuovo M, Fiorino G, Gatta E, Robello M, Borghi R, Tabaton M, Russo C, Dieci G, Cancedda R, Pagano A (2011) RNA polymerase III drives alternative splicing of the potassium channel-interacting protein contributing to brain complexity and neurodegeneration. J Cell Biol 193: 851-66

Meacham GC, Patterson C, Zhang W, Younger JM, Cyr DM (2001) The Hsc70 co-chaperone CHIP targets immature CFTR for proteasomal degradation. Nat Cell Biol 3: 100-5

Monaghan MM, Menegola M, Vacher H, Rhodes KJ, Trimmer JS (2008) Altered expression and localization of hippocampal A-type potassium channel subunits in the pilocarpine-induced model of temporal lobe epilepsy. Neuroscience 156: 550-62

Mycko MP, Cwiklinska H, Walczak A, Libert C, Raine CS, Selmaj KW (2008) A heat shock protein gene (Hsp70.1) is critically involved in the generation of the immune response to myelin antigen. Eur J Immunol 38: 1999-2013

Nadal MS, Ozaita A, Amarillo Y, Vega-Saenz de Miera E, Ma Y, Mo W, Goldberg EM, Misumi Y, Ikehara Y, Neubert TA, Rudy B (2003) The CD26-related dipeptidyl aminopeptidase-like protein DPPX is a critical component of neuronal A-type K+channels. Neuron 37: 449-61

Norris AJ, Nerbonne JM (2010) Molecular dissection of I(A) in cortical pyramidal neurons reveals three distinct components encoded by Kv4.2, Kv4.3, and Kv1.4 alphα-subunits. J Neurosci 30: 5092-101

Racine RJ (1972) Modification of seizure activity by electrical stimulation. II. Motor seizure. Electroencephalogr Clin Neurophysiol 32: 281-94

Rejdak K, Kuhle J, Ruegg S, Lindberg RL, Petzold A, Sulejczak D, Papuc E, Rejdak R, Stelmasiak Z, Grieb P (2012) Neurofilament heavy chain and heat shock protein 70 as markers of seizure-related brain injury. Epilepsia 53: 922-7

Schlecht R, Scholz SR, Dahmen H, Wegener A, Sirrenberg C, Musil D, Bomke J, Eggenweiler HM, Mayer MP, Bukau B (2013) Functional analysis of Hsp70 inhibitors. PLoS One 8: e78443

Shibata R, Nakahira K, Shibasaki K, Wakazono Y, Imoto K, Ikenaka K (2000) A-type K+ current mediated by the Kv4 channel regulates the generation of action potential in developing cerebellar granule cells. J Neurosci 20: 4145-55

Singh B, Ogiwara I, Kaneda M, Tokonami N, Mazaki E, Baba K, Matsuda K, Inoue Y, Yamakawa K (2006) A Kv4.2 truncation mutation in a patient with temporal lobe epilepsy. Neurobiol Dis 24: 245-53

Soh H, Goldstein SA (2008) I SA channel complexes include four subunits each of DPP6 and Kv4.2. J Biol Chem 283: 15072-7

Tang YQ, Liang P, Zhou J, Lu Y, Lei L, Bian X, Wang K (2013) Auxiliary KChIP4a suppresses A-type K+ current through endoplasmic reticulum (ER) retention and promoting closed-state inactivation of Kv4 channels. J Biol Chem 288: 14727-41

Tang YQ, Zhou JH, Yang F, Zheng J, Wang K (2014) The tetramerization domain potentiates Kv4 channel function by suppressing closed-state inactivation. Biophys J 107: 1090-104

Tsaur ML, Sheng M, Lowenstein DH, Jan YN, Jan LY (1992) Differential expression of K+ channel mRNAs in the rat brain and down-regulation in the hippocampus following seizures. Neuron 8: 1055-67

Walker VE, Wong MJ, Atanasiu R, Hantouche C, Young JC, Shrier A (2010) Hsp40 chaperones promote degradation of the HERG potassium channel. J Biol Chem 285: 3319-29

Wang AM, Miyata Y, Klinedinst S, Peng HM, Chua JP, Komiyama T, Li X, Morishima Y, Merry DE, Pratt WB, Osawa Y, Collins CA, Gestwicki JE, Lieberman AP (2013) Activation of Hsp70 reduces neurotoxicity by promoting polyglutamine protein degradation. Nat Chem Biol 9: 112-8

White AM, Williams PA, Ferraro DJ, Clark S, Kadam SD, Dudek FE, Staley KJ (2006) Efficient unsupervised algorithms for the detection of seizures in continuous EEG recordings from rats after brain injury. J Neurosci Methods 152: 255-66

Williamson DS, Borgognoni J, Clay A, Daniels Z, Dokurno P, Drysdale MJ, Foloppe N, Francis GL, Graham CJ, Howes R, Macias AT, Murray JB, Parsons R, Shaw T, Surgenor AE, Terry L, Wang Y, Wood M, Massey AJ (2009) Novel adenosine-derived inhibitors of 70 kDa heat shock protein, discovered through structure-based design. J Med Chem 52: 1510-3

Yang T, Hsu C, Liao W, Chuang JS (2008) Heat shock protein 70 expression in epilepsy suggests stress rather than protection. Acta Neuropathol 115: 219-30

Yenari MA, Fink SL, Sun GH, Chang LK, Patel MK, Kunis DM, Onley D, Ho DY, Sapolsky RM, Steinberg GK (1998) Gene therapy with HSP72 is neuroprotective in rat models of stroke and epilepsy. Ann Neurol 44: 584-91

Young JC (2014) The role of the cytosolic HSP70 chaperone system in diseases caused by misfolding and aberrant trafficking of ion channels. Dis Model Mech 7: 319-29

Zagha E, Ozaita A, Chang SY, Nadal MS, Lin U, Saganich MJ, McCormack T, Akinsanya KO, Qi SY, Rudy B (2005) DPP10 modulates Kv4-mediated A-type potassium channels. J Biol Chem 280: 18853-61

Zhang Y, Nijbroek G, Sullivan ML, McCracken AA, Watkins SC, Michaelis S, Brodsky JL (2001) Hsp70 molecular chaperone facilitates endoplasmic reticulum-associated protein degradation of cystic fibrosis transmembrane conductance regulator in yeast. Mol Biol Cell 12: 1303-14

Zhou J, Tang Y, Zheng Q, Li M, Yuan T, Chen L, Huang Z, Wang K (2015) Different KChIPs compete for heteromultimeric assembly with pore-forming Kv4 subunits. Biophys J 108: 2658-69

